# Scalable De Novo Classification of Antibiotic Resistance of Mycobacterium Tuberculosis

**DOI:** 10.1101/2023.11.16.567394

**Authors:** Mohammadali Serajian, Simone Marini, Jarno N. Alanko, Noelle R. Noyes, Mattia Prosperi, Christina Boucher

## Abstract

We develop a robust machine learning classifier using both linear and nonlinear models (i.e., LASSO logistic regression (LR) and random forests (RF)) to predict the phenotypic resistance of *Mycobacterium tuberculosis* (MTB) for a broad range of antibiotic drugs. We use data from the CRyPTIC consortium to train our classifier, which consists of whole genome sequencing and antibiotic susceptibility testing (AST) phenotypic data for 13 different antibiotics. To train our model, we assemble the sequence data into genomic contigs, identify all unique 31-mers in the set of contigs, and build a feature matrix *M*, where *M* [*i, j*] is equal to the number of times the *i*-th 31-mer occurs in the *j*-th genome. Due to the size of this feature matrix (over 350 million unique 31-mers), we build and use a sparse matrix representation. Our method, which we refer to as MTB++, leverages compact data structures and iterative methods to allow for the screening of all the 31-mers in the development of both LASSO LR and RF. MTB++ is able to achieve high discrimination (F-1 greater than 80%) for the first-line antibiotics. Moreover, MTB++ had the highest F-1 score in all but three classes and was the most comprehensive since it had a F-1 score greater than 75% in all but four (rare) antibiotic drugs. We use our feature selection to contextualize the 31-mers that are used for the prediction of phenotypic resistance, leading to some insights about sequence similarity to genes in MEGARes. Lastly, we give an estimate of the amount of data that is needed in order to provide accurate predictions.

## INTRODUCTION

The World Health Organization (WHO) estimates that there were over 10 million cases of tuberculosis worldwide in 2019, resulting in over 1.4 million deaths, with a worrisome increasing trend yearly. The disease is usually caused by *Mycobacterium tuberculosis* (MTB), through airborne transmission. Treatment of tuberculosis is estimated to be 85% successful, however, this percentage drops to 57% with MTB that exhibits resistance to multiple antibiotic drugs [35], for which fewer treatment options are available. Thus, the identification of antibiotic resistance is critical for patient care management, guiding antibiotic drug stewardship, and decreasing the selection of resistant strains from a public health and ecological standpoint. Typically, antibiotic resistance is measured through the Antibiotic Susceptibility Test (AST) for each antibiotic resistant drug. DNA sequencing can be paired with AST to identify genetic differences between antibiotic resistant and susceptible strains, both in clinical and ecological settings. Several data and model resources for MTB have been established to support research on antibiotic resistance. One of the largest is the international Comprehensive Resistance Prediction for Tuberculosis: an International Consortium (CRyPTIC) Consortium [34], which performed whole genome sequencing for tens of thousands of MTB isolates collected from diverse locations worldwide as well as AST analysis of these isolates for over a dozen antibiotic drugs.

Recently, the CRyPTIC database has been used for a genome-wide association study to identify oligopeptides and oligonucleotides in MTB associated with resistance to single antibiotics [33, 34]. These findings confirmed and expanded on prior work that identified single nucleotide polymorphisms (SNPs) that are associated with antibiotic drug resistance in MTB [29–31]. Prior methods for classifying antibiotic resistance in MTB use catalogues of genetic variants, i.e., predominantly SNPs: TBProfiler [27], PhyResSE [13], and KvarQ [32]. TBProfiler [27] combine several tools including Trimmomatic [4], BWA [24] or Bowtie 2 [22], BCFtools, and SAMtools [11], to map sequence reads from whole genome sequencing to a WHO-endorsed catalogue of 17,356 MTB variants whose resistance is known. Similarly, PhyResSE [13] is a pipeline that also uses several third-party methods, including QualiMap [15], SAMtools [25], and GATK [26]. Unlike TBProfiler, PhyResSE aims to identify both the lineage and resistance type from whole genome sequence data. PhyResSE aligns all the input reads to the MTB H37Rv reference genome using BWA [23]; identifies variants using GATK [26]; and predicts the resistance and lineage based on the found variants. KvarQ [32] also uses a catalogue of known MTB variants to identify antibiotic resistance, but rather than aligning all reads to a database to detect variants, it extracts the relevant information from each individual read. One of the main contributions of KvarQ was eliminating the need for alignment or assembly of all the reads.

In addition to these, there are other methods of pairing machine learning or combinatorial approaches with variant catalogues to identify antibiotic resistance in MTB. These include Mykrobe [7, 20], GenTB [21], and the models by Kuang et al. [21]. Mykrobe, which was initially released in 2015 [7] and then updated in 2019 [20], relies on the construction and analysis of a de Bruijn graph from catalogues of resistant and susceptible alleles on different genetic backgrounds, along with a set of antibiotic resistance genes. Together, they form what the authors refer to as a *reference graph*. The reference graph is compared to the de Bruijn graph of the sequence data and through this comparison, a prediction of the resistance is made based on statistical tests. In 2019, Mykrobe was updated to include a mutation catalogue, giving greater sensitivity to detect pyrazinamide resistance; another improvement was to allow for a user-specified catalogue. In 2021, GenTB [17] was released. GenTB is a machine-learning method that classifies resistance to 13 different drugs using a catalogue of variant positions spanning 18 resistance-associated genetic loci. It trains both a random forest classifier and a deep neural network. A more recent work was released by Kuang et al. [21] in 2022, training concurring machine learning models (LR, RF, and convolutional neural network) using a dataset of 10,575 MTB isolates for which both the lineage and AST results are known.

Lastly, there are general-purpose methods for classifying antibiotic resistance from either whole genome sequence data or metagenomic sequence data. Most of these methods and databases, including MEGARes [5], contain curated antibiotic genes for *MTB* resistance, but have not been specifically evaluated on MTB isolates. The exception to this is ResFinder [6, 14], a generally purposed method that stands out as having been redeveloped and evaluated for MTB phenotypic resistance classification.

In this paper, we develop a robust machine-learning classifier that predicts the phenotypic resistance of MTB for a broad range of antibiotic drugs. Our method, which we refer to as MTB++, classifies the resistance to 16 different antibiotics of an isolate based on the oligonucleotides in its whole genome sequence data. To the best of our knowledge, this is the first machine learning method that is trained in a *de novo* fashion, meaning that it only uses the oligonucleotides and AST data for training; whereas existing classifiers—including ResFinder [6, 14], GenTB [21], TBProfiler [27], Mykrobe [7], and KvarQ [32]—require prior knowledge of genetic variants that associate with MTB antibiotic resistance occurrence. It should be noted that the recent CRyPTIC study [33] also considered oligonucleotides without confining interest to genetic variants, but this analysis did not include the release of a classifier. Moreover, in order to construct our classifier we consider up to three orders of magnitude more oligonucleotides in comparison to the CRyPTIC study [33]. We enable the training of this large search space through a combination of succinct data structures and machine learning. The combination of our consideration of a large search space and a *de novo* machine learning approach is that it allows for novel mechanisms of resistance to be identified and for increased performance accuracy to be obtained. We evaluated our method against all available competing methods. MTB++ is able to achieve high discrimination (i.e., F-1 score over 80%) for 10 out of the 13 antibiotic drugs we considered that had a data balance (ratio of isolates with phenotypic resistance to total number of isolates) of over 5%. Competing methods were only able to achieve high discrimination for five to seven of these antibiotic drugs. In addition to considering the accuracy of the classification, we identify antibiotic resistance genes containing the oligonucleotides of our models to hypothesize about novel variants associated with resistance in MTB. The trained classifier, and source code to create the classifier are publicly available on GitHub at https://github.com/M-Serajian/MTB-Pipeline.

## RESULTS

### MTB++ was the most comprehensive classifier, generating accurate predictions on the largest number of antibiotic drugs

We give an overview of our methods in Figure 1. We downloaded all publicly available whole genome sequencing data, and phenotypic data for MTB from the CRyPTIC Consortium [33]. We give the number of isolates that were deemed to be susceptible, resistant, and ambiguous to each antibiotic drug based on the CRyPTIC AST in Table **??**. The percentage that is resistant is given since it has a significant effect on the performance of the methods we evaluated. We then assembled the sequence data from all isolates in CRyPTIC that have both phenotypic and genomic data and identified the set of all unique 31-mers from these contigs extracted as features for classification. This resulted in 6,224 sets of contigs, 350 million 31-mers before filtering and close to 17 million 31-mers after filtering. The data was then split into training and testing data and both a LR model and a RF model were trained and used in order to achieve optimal performance. Five-fold cross-validation was used for evaluation. We report the F-1 scores in Table 1. In addition to the F-1 score, we illustrate the Receiver Operating Characteristic (ROC) curve for each antibiotic drug in figure 4 in the Supplement. Our method achieved an F-1 score greater than 90% in four classes, and another eight classes had an F-1 score greater than 75%.

**Table 1:**
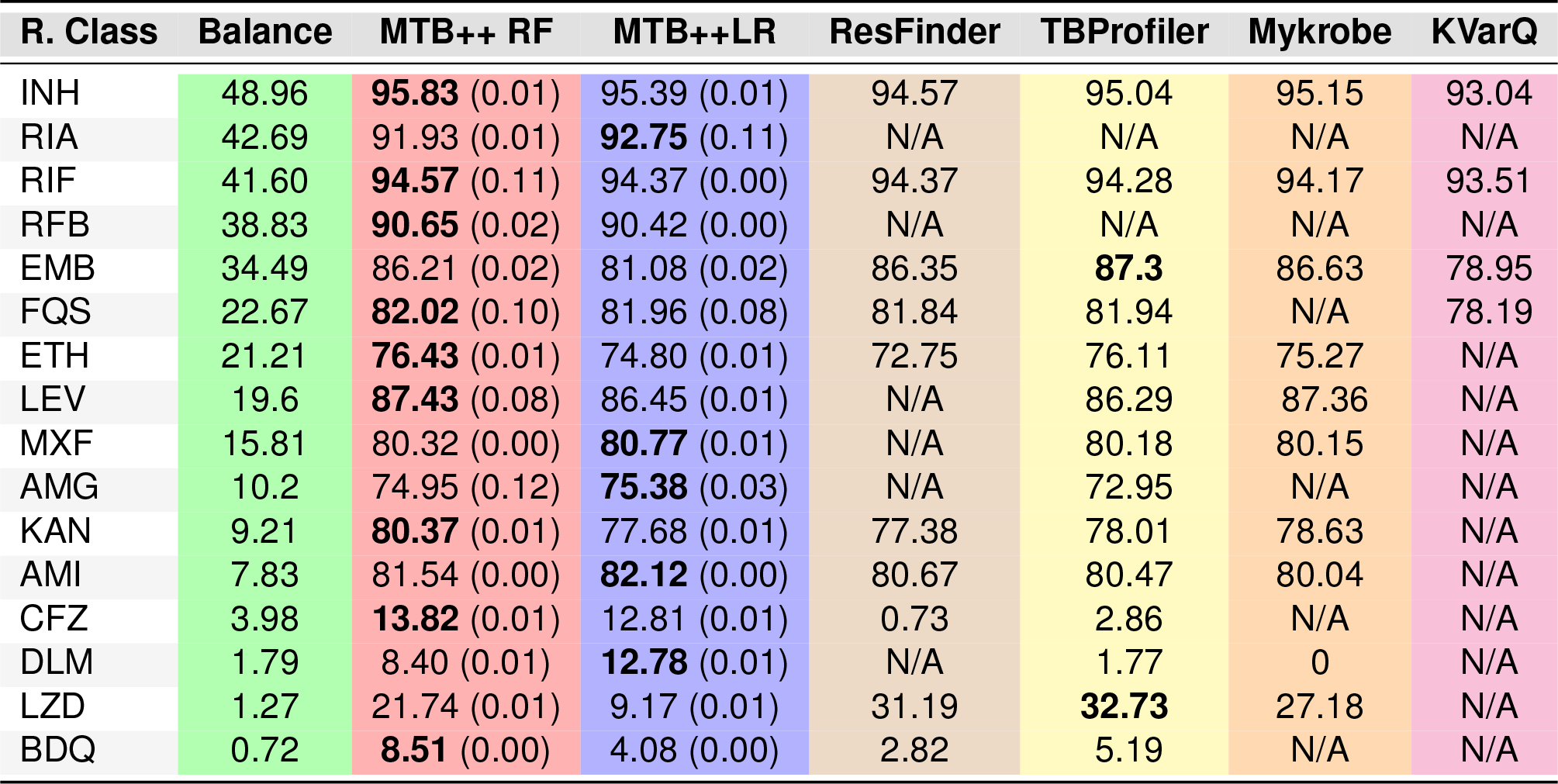
Comparison between MTB++’s random forest (RF) and linear regression (LR) with other methods. Our performance is demonstrated by the mean and (std) of the F-1 score in cross-validation. For some of the competing methods, some of our drugs in this study were not covered, and those cases are demonstrated using N/A.

**Figure 1.**
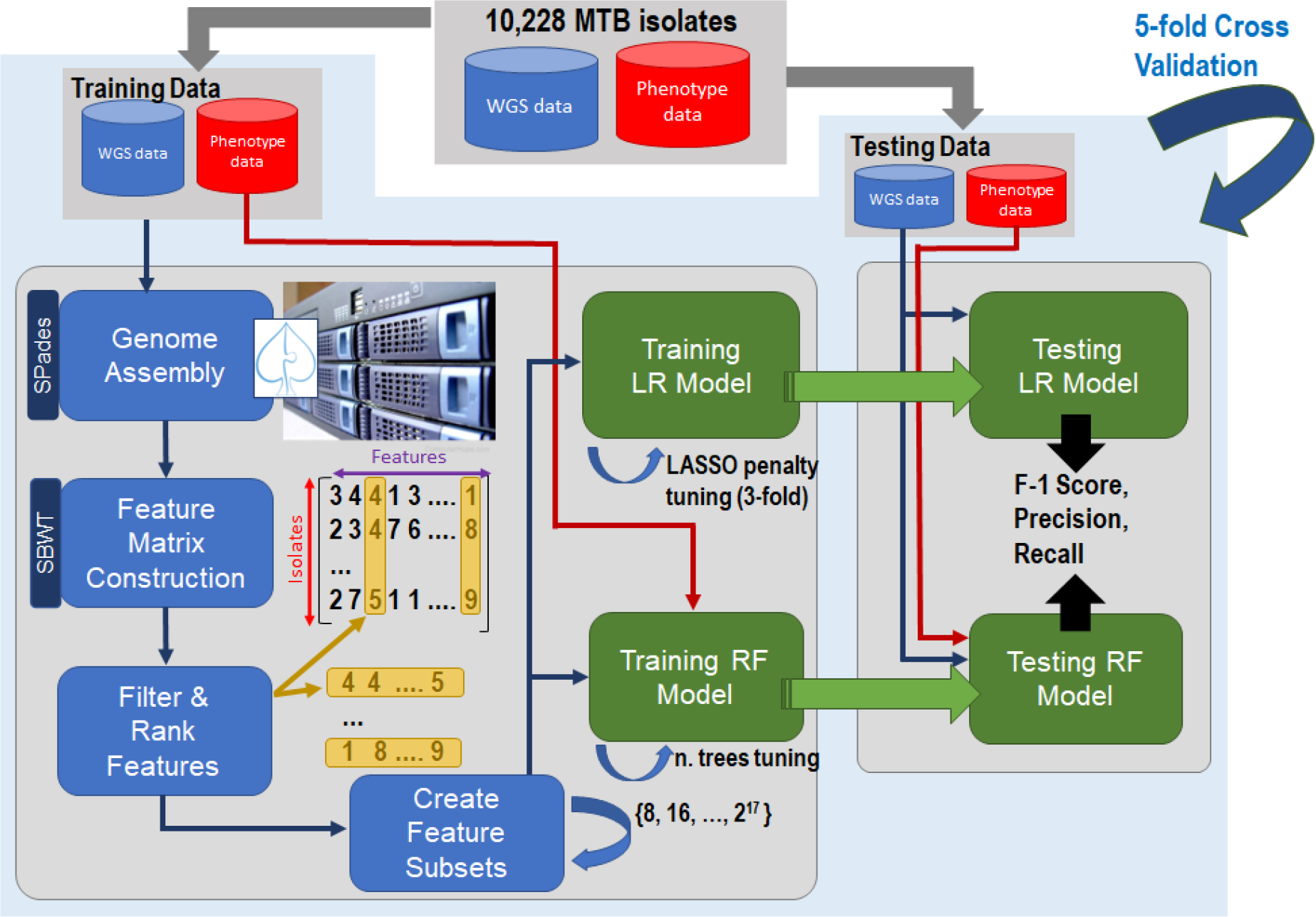
Overview of MTB++. We downloaded all publicly available data available through CRyP-TIC [34]. Split the data into training and testing datasets. We trained both a Linear Regression (LR) and Random Forest (RF) model on each dataset. The training data was assembled using SPAdes [2], then all unique 31-mer were extracted in order to create a feature matrix for training the models. Five-fold cross-validation was used to evaluate the trained models and calculate the F-1 score.

We compared the performance of MTB++, ResFinder [6, 14], Mykrobe [20], KvarQ [32], and TBProfiler [8, 27]. We note that we were unable to compare against PhyResSE[13], GenTB [17], and the method of Kuang et al. [21]^1^. MTB++ had the highest F-1 score for 13 of the 16 antibiotic drugs evaluated and achieved a performance of greater than 90% for two antibiotics (rifamycin and rifabutin)in which the competing methods were unable to make any prediction. KVarQ was only able to perform a prediction for four out of the 16 antibiotic drugs. Mykrobe was able to achieve performance for nine of the 16 antibiotics but had lower performance than MTB++ in eight of those nine antibiotics. Mykrobe outperformed MTB++ for linezolid, yet the F-1 score of both methods was less than 28%. TBProfiler and ResFinder were the most competitive but ResFinder only had acceptable performance (i.e., greater than 70 %) in seven of 16 antibiotics. TBProfiler achieved acceptable performance on 10 out of the 16 antibiotics but had a lower F-1 score than MTB++ on nine out of those 10 antibiotics. In summary, MTB++ had superior performance than all competing methods for almost all antibiotic drugs; the only ones for which the competing methods outperformed our method were ethambutol and rare antibiotic drugs which have highly unbalanced data (the ratio of isolates with resistant phenotype to total number of isolates is below 40%). Moreover, all methods were unable to achieve acceptable performance on four of the 16 antibiotics, where the data were highly unbalanced; these antibiotics are bedaquiline, linezolid, delamanid, and clofazimine. Each of these antibiotic drugs is reserved for the use of multi-drug-resistant MTB in order to ensure judicious use and because of their numerous side effects, including drug interactions and even death as in the case of bedaquiline [10].

### Ranking of features reveals genomic sequences that have a high association with specific classes of resistance

One of the advantages of MTB++ is that it is *de novo*, meaning that it uses no prior knowledge about the genetic variations that have been confirmed to be associated with MTB antibiotic resistance. Our method begins by considering all unique 31-mers, resulting in over 350 million 31-mers and filters for those that occur sparingly (less than 10 times) or ubiquitously (more than 3,000 times). From the set of filtered 31-mers, we trained the models in an exponentially larger set of 31-mers, i.e., from 2^8^ 31-mers to 2^17^ 31-mers, and reported the performance of the model that obtained the maximum F-1 score. The number of features that had a non-zero coefficient from these models is reported in Table 2. These features were then ranked using the chi-square test.

This analysis of features demonstrates high redundancy between 31-mers. In particular, the application of LASSO regularization in the training of a LR model, resulting in a substantial reduction in the number of features with non-zero coefficients relative to the total feature count, offers valuable insights into the underlying extracted 31-mers. This phenomenon underscores the presence of feature redundancy and inter-feature correlations within the 31-mers for each antibiotic drug. Feature redundancy occurs when multiple 31-mers convey similar or redundant information (belonging to the same mechanism of resistance), leading LASSO to select one representative feature while penalizing others. This selective process aids in feature dimensionality reduction and model simplification. In addition, the presence of inter-feature correlation implies that some features are highly correlated with others, making them less individually informative. LASSO, in its feature selection process, tends to favor one feature from a correlated group, effectively pruning the others. This not only enhances model interpretability but also reduces (multi-)collinearity that is caused by the correlations between features, which can destabilize coefficient estimates and hinder the generalizability of the model.

Moreover, after training the RF models, a sudden decrease in the significance (Gini impurity measure) of 31-mers can be noticed for all antibiotics. In this context, most features are classified as unimportant while only a few of the 31-mers are deemed significant, which can be attributed to the ensemble nature of the RF algorithm and the inherent feature selection process it employs. RF algorithms construct numerous decision trees on bootstrapped data subsets and aggregate their predictions, leading uninformative or redundant features to consistently rank lower in importance across the ensemble. Collinearity or multicollinearity of the features causes the RF to distribute importance scores among them, thus diminishing their individual significance. Consequently, a less informative, uncorrelated feature may receive a higher importance score as it contributes unique information to the model. This fact underscores the importance of considering both feature importance and feature correlations when interpreting RF results, ensuring a nuanced understanding of the model’s behavior. Therefore, to reduce the negative impact of collinearity to identify the most informative 31-mers, we combined the top features selected by both RF and LR as the most informative 31-mers.

### Contextualizing the features leads to model validation and association between antibiotic drugs

In order to further evaluate and use the model, we extracted all features that had a non-zero coefficient from the model that obtained the optimal F-1 score for each antibiotic drug, and aligned these features to MEGARes 3.0 [5] database using BWA [24] with the restriction that we only considered exact matches. MEGARes has over 8,000 AMR genes with less than two dozen of these being MTB-specific antibiotic resistance genes. MEGARes also contain metal resistance and virulence factors. After alignment, we omitted any AMR genes where there were less than ten 31-mers aligned to it; this was done to remove any spurious results. Tables 4 and 5 give the MEGARes genes by antibiotic drugs for each MEGARes resistance class. Table 4 is restricted to antibiotic drugs for which the balance of the data is at least 15%. Table 5 is restricted to antibiotic drugs for which the balance of the data is at most 11%. In order to further contextualize our results, we first considered the possible multi-drug resistance of the isolates. Figure 2a illustrates the frequency of resistance to two different antibiotic drugs. Complementary to this, we also calculated the Jaccard similarity index of the set of features for each pair of antibiotic drugs. The Jaccard similarity compares members for two sets to determine which members are shared and which are distinct. Figure 2b gives an illustration of Jaccard similarity.

**Figure 2.**
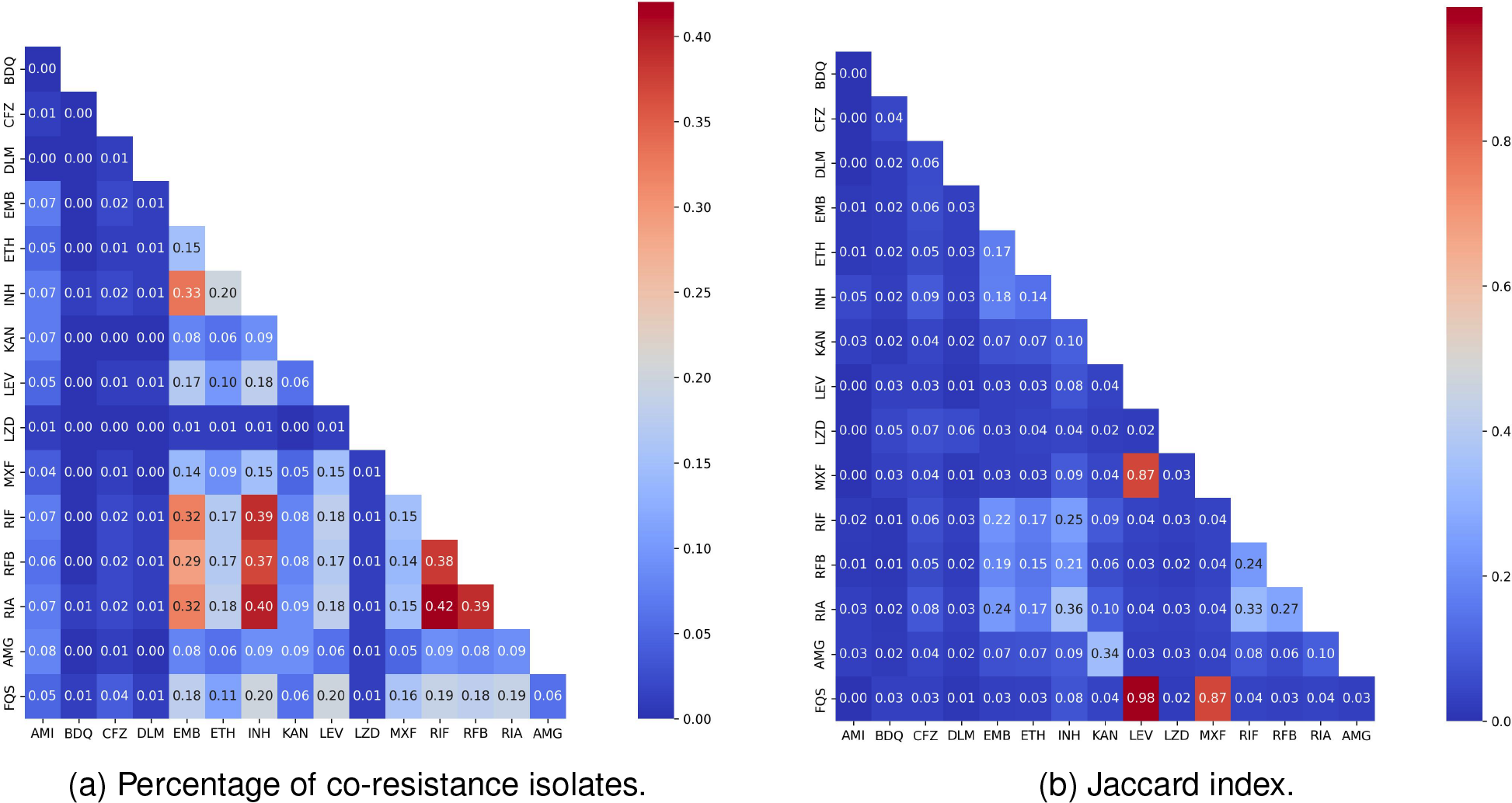
Figure (a) is a heatmap illustrating the Jaccard index for shared 31-mers between all pairs of antibiotic drugs, normalized by the total number of 31-mers in the union of features for the two antibiotic drugs. Figure (b) is a heatmap that gives the ratio of isolates resistant to two distinct antibiotic drugs to the total number of isolates.

Figure 2a illustrates that there is high frequency of co-resistance between rifamycin and ri-fampicin (42%), rifamycin and rifabutin (39%), and rifampicin and rifabutin (38%). It is worth noting that these antibiotic drugs are associated with a larger family of resistance: rifamycins, which is a first-line therapy for mycobacterial infections. Isoniazid resistance was also associated with resistance to rifamycin, rifampicin, and rifabutin. However, there was not as strong of a correlation with the Jaccard index between these classes of resistance as can be seen in Figure 2b. Our results in Table 4 are reflective of this similarity. Isoniazid, rifamycin, rifampicin, and rifabutin had exact alignments to several of the same genes in MEGARes, including MEG 8171 (katG), MEG 6144 (rpsL), MEG 3237 (GyrA), MEG 2710, MEG 2711 (embB), MEG 7259 (remB), MEG 6090, MEG 6134 (rpoB). Rifabutin is uniquely associated with MEG 3180 (GyrA), which is associated with fluoroquinolone resistance [36].

Other pairs of antibiotic drugs that had high Jaccard similarity were: levofloxacin-fluoroquinolone, moxifloxacin-fluoroquinolone, and moxifloxacin-levofloxacin. Levofloxacin and moxifloxacin are frequently classified as fluoroquinolones, which validates our model. This relationship is further reflected in Table 4 as MEG 2337, which is a fluoroquinolone antibiotic resistance gene, has the largest number of exact alignments to levofloxacin, fluoroquinolone, and moxifloxacin.

There were several AMR genes that were unique to specific antibiotic drugs. These associations validate our model and demonstrate the possible novel mechanisms of resistance. MEG 2712 was uniquely associated with ethambutol. This AMR gene is defined as being in the ethambutol class in MEGARes, and targets arabinosyltransferase. Hence, this result validates our findings. More surprisingly, ethambutol was uniquely associated with MEG 2653 and MEG 2653, which corresponds to copper resistance in MEGARes. MEG 8079, which is classified as a MTB-specific AMR gene in MEGARes, also targets arabinosyltransferase. This gene was uniquely associated to rifampicin. Interestingly ethambutol and rifampicin are frequently combined for the treatment of MTB so it is conceivable that there is a region in MEG 8079 that is encoding for rifampicin resistance or this was due to multi-drug resistance caused by the concurrent treatment with ethambutol and rifampicin. Lastly, ethambutol was also associated with MEG 1490, which is defined as a cationic antibiotic peptide (CAMP) resistance gene in MEGARes.

### MTB++ reveals genomic features associated with rare antibiotic drugs and highlights the need for more data for “last resort” antibiotic drugs

As shown in Table 1, we witness that the performance of MTB++ was highly dependent on the balance of the data. MTB++ had an F-1 score of over 90% for antibiotic drugs that had balanced data, i.e., greater than 38%. When the balance was below 5% all the models struggled with adequate performance; for clofazimine, delamanid, linezolid, and bedaquiline all the models had a F-1 score of 33 % or below. Next, we went beyond the balance of the data and considered the number of isolates that were deemed to be phenotypically resistant in the CRyPTIC dataset. Hence, Figure 3 illustrates the number of isolates versus the F-1 score for each antibiotic drug. When the number of resistant isolates was at least 500 there was a sharp increase in the performance of both the RF and LR models. In the case of antibiotic drugs representing resistance to antibiotics prescribed as a last resort (i.e., bedaquiline and linezolid) that had a balance of less than 1.3% (or less than 80 resistant isolates), the performance was degraded as the model struggled to find enough significant features to distinguish the resistant isolates. Notwithstanding, it should be noted that even for rare antibiotic drugs—where the balance is less than 5%—our model achieved a F-1 score between 8% and 22%. This implies that our model is finding oligonucleotides that are associated with the various antibiotic drugs and could be related to new and/or undiscovered mutations that are not among currently known variants.

**Figure 3.**
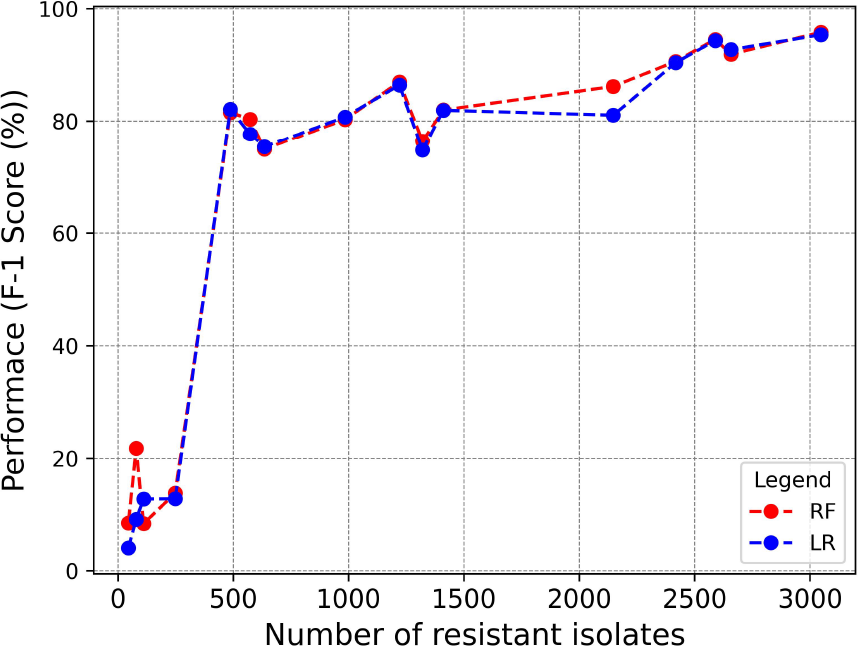
Illustration of the relationship between the optimal F-1 score (y-axis) and the number of resistant isolates (x-axis).

**Figure 4.**
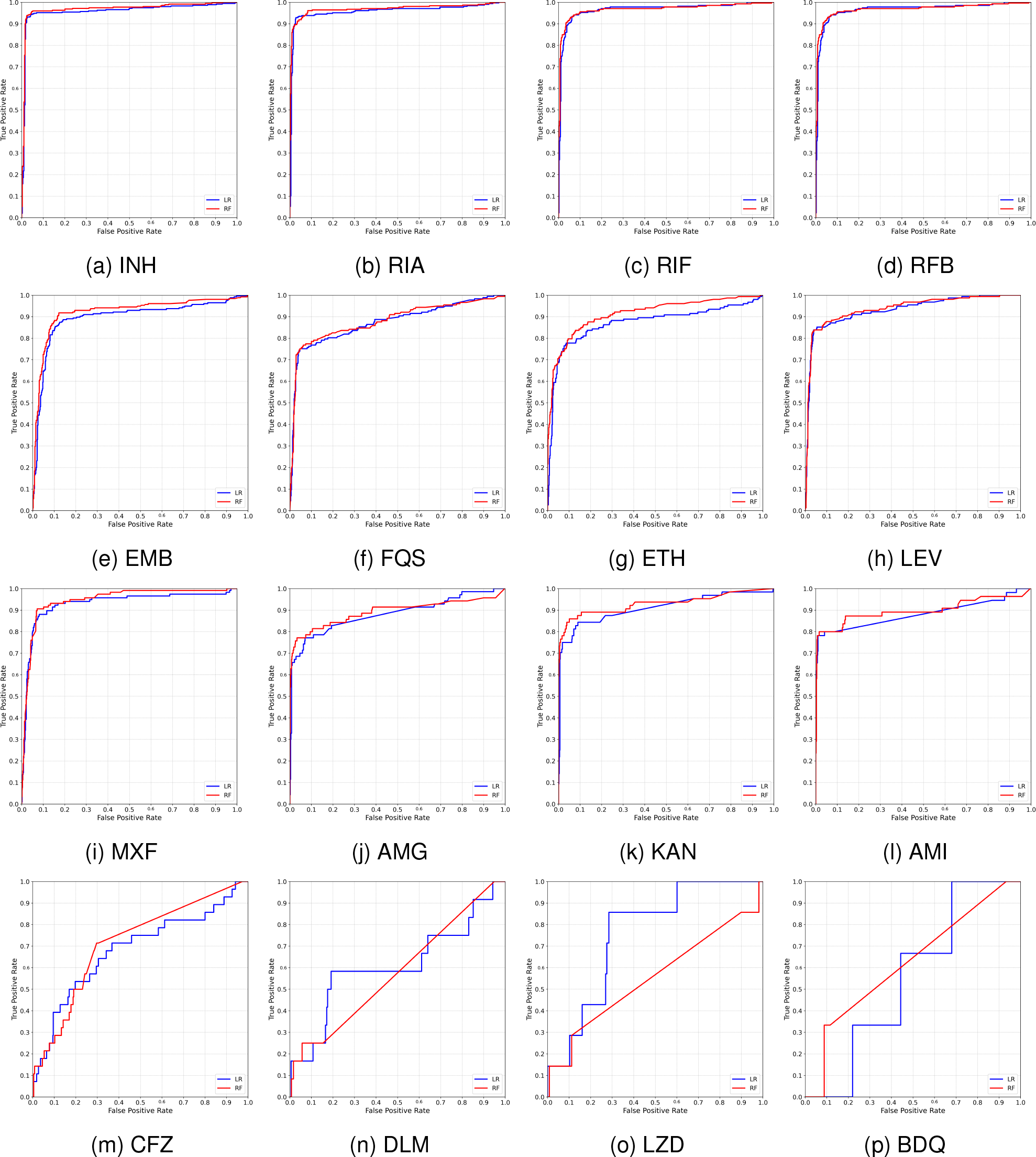
ROC curve of the classification models for each resistance phenotype. The red color is related to RF and the green color is related to LR. The best LR and RF for each drug are highlight by dark red and green.

From Table 5, we show that amikacin, kanamycin, and aminoglycoside have a considerable number of common antibiotic resistance genes and that most of these antibiotic resistance genes belong to the aminoglycoside class in MEGARes. MEG 5 was uniquely found for these three antibiotic drugs. Amikacin and kanamycin belong to the same family of antibiotics known as aminoglycosides so this association helps validate our model.

Delamanid and linezolid had a very low number of phenotypically resistant isolates, and thus, had a data balance of 1.79% and 1.27%, respectively; only bedaquiline had a lower balance (i.e., 0.72%). Delamanid had an association to a multi-drug antibiotic resistance gene (MTRAD), and a biocide resistance antibiotic resistance gene (MDTK). Further investigation into these associations is warranted but appears to be reasonable given that these antibiotics are considered last-resort drugs used for multi-drug resistant MTB. Figures 2b and 2a demonstrate that very few isolates displayed co-resistance with delamanid. Moreover, there were no other antibiotic drugs that had a Jaccord index of more than 0.03 with delamanid.

Linezolid resistance was associated with MEG 1490, an antibiotic resistance gene linked to CAMP resistance that was also associated with resistance to ethionamide. It should be noted that there were very few isolates that had phenotypic resistance to both ethionamide and linezolid; as shown in Figure 2a. Phenotypic resistance to linezolid was uniquely associated with the oxazolidinone antibiotic resistance class in MEGARes (MEG 8670). Oxazolidinones, a new chemical class of synthetic antibiotics, has a unique mechanism of action that involves the inhibition of bacterial protein synthesis. Linezolid drugs belong to the oxazolidinone class of antibiotics [19] so this finding appears to also validate the model.

## DISCUSSION

We presented a trained classifier that is able to accurately predict the resistance profile for 12 different antibiotic drugs, including some rare antibiotic drugs that are used for highly-resistant forms of MTB. This suggests that the genetic features for these antibiotics are well conserved and able to used for classification. The performance of MTB++ suggests that accurate performance of antibiotic resistance of MTB can be achieved for a wider range of antibiotic drugs. This is critically important since competing methods make accurate predictions for a more limited set of antibiotic drugs. We note that one of the main advantages of our method is that its prediction uses no prior knowledge of the genetic resistance-associated variants. More specifically, MTB++ is trained using all unique 31-mers from assembled whole genome sequence data along with side-by-side AST data describing the phenotypic resistance. This *de novo* construction of the feature set requires us to consider three orders of magnitude more features than prior studies before filtering and over two times more features after filtering—inclusive in this comparison is the genome-wide association study from the CRyPTIC Consortium [33] that uses the same dataset but limits interest to approximately five million 31-mers. We note that after filtering, the feature matrix is over 1.2 terabytes in size. The space- and memory-efficient method of Alanko et al. [1] for building and storing the feature matrix allows us to build, train, and analyze a classifier on a feature matrix of this magnitude. In particular, the alignment of 31-mers to MEGARes bridges MTB resistance prediction with AMR database creation, and is possible due through the application of succinct data structures. This area of combining succinct data structures with machine learning classification is still in a nascent stage but as biological datasets increase in size, we believe this will quickly become an area that requires further exploration.

## METHODS

### Dataset Description

We downloaded all publicly available whole genome sequencing data, and AST data for MTB from the CRyPTIC Consortium [33]. The data are available on the European Nucleotide Archive (ENA) FTP server. The whole genome sequencing data are all Illumina paired-end reads and are annotated using ERR numbers. Resistance phenotype data are annotated with ERS numbers.

We matched the ERR and ERS for each isolate using a .JSON file. The ERS numbers and JSON file are available here: https://ftp.ebi.ac.uk/pub/databases/cryptic/release_june2022/ We note that isolates were removed from further consideration if either the sequence data could not be retrieved or the ERR and ERS number could not be properly matched. After the removal process, we obtained a total of 6,224 isolates with complete sequence.

Based on the MIC threshold used by the CRyPTIC consortium (Table 3), we labeled each isolate as resistant or susceptible to 13 antibiotic drugs: amikacin (AMI), bedaquiline (BDQ), clofazimine (CFZ), delamanid (DLM), ethionamide (ETH), ethambutol (EMB), isoniazid (INH), kanamycin (KAN), levofloxacin (LEV), linezolid (LZD), moxifloxacin (MXF), rifampicin (RIF) and rifabutin (RFB). In the treatment of TB, first-line antibiotic drugs consist of EMB, INH, and RIF; second-line drugs are AMI, ETH, KAN (injectable agent), LEV, MXF, and RFB; and the last line of new and repurposed drugs are BDQ, CFZ, DLM, and LZD [33]. The number of isolates resistant to the last line of antibiotic drugs was very low in comparison to the first two lines (e.g., less than 1% for BDQ). Thus, we combined some of these drugs into classes based on the classification hierarchy of Doster et al. [12]: (1) KAN and AMI were combined into the aminoglycoside (AMG) group, (2) RFB and RIF were combined into the rifamycin (RIA) group, and (3) MXF, LEV and CFZ were combined into the fluoroquinolones (FQS) group. The combination process is as follows: if an isolate is phenotypically resistant to at least one of the antibiotic drugs in a group, then that isolate will be considered as resistant to that group. In Table **??**, we provide the number of isolates that were phenotypically susceptible, ambiguous, and resistant isolates, as well as, the proportion of isolates that were resistant to each antibiotic out of the total number of isolates (excluding ambiguous). The phenotypic resistance for two of the antibiotic drugs, ethambutol, and ethionamide, was “intermediate”; in those cases, we labeled the isolates as phenotypically resistant since they still show a resistant phenotype.

### Genome Assembly and Feature Extraction

We assembled the sequence reads for all 6,224 isolates using SPAdes (version 3.15.3) [28], and evaluated the quality of the assemblies using Quast [18]. The mean and standard deviation (std) for the number of reads for each isolate is 4,397,842 and 2,685,471, respectively. The mean and standard deviation of the N50 is 96,971.55 and 44,104.37, respectively.

Next, we used all unique 31-mers from the assembled genomes as the features for our machine learning models. Thus, we created an *n* × *m* integer matrix that stores the number of occurrences of each 31-mer in each assembled genome, where *n* is the number of unique 31-mers in all the assembled genomes, and *m* is the number of genomes. Hence, the integer at the *i*-th row and *j*-th column are equal to the number of times the *i*-th 31-mer occurs in the *j*-th assembled genome. To build this matrix, we first constructed a perfect hash function that maps the unique 31-mers of the data to the rows of the color matrix. We used the SBWT data structure of Alanko et al. [1] to implement this hash function. The hash value of 31-mer *x* is the number of distinct 31-mers in the data that are co-lexicographically smaller than *x*. For our data, the matrix has 356,359,267 rows (31-mers, *n*) and 6,224 columns (genomes, *m*). Since the matrix is very large, we only store the non-zero elements to save space (ASCII format). That is, for each row in the matrix, we store a list of pairs (*g*_1_, *c*_1_), (*g*_2_, *c*_2_), …, where pair (*g*_i_, *c*_i_) indicates that genome *g*_i_ contains *c*_i_ copies of the 31-mer of the row. These lists are built by iterating the columns (genomes) left to right and using the hash function to look up the rows of the 31-mers, creating new counter pairs (*g*_i_, *c*_i_) on demand when a *k*-mer is seen the first time in the genome, and incrementing counters in existing pairs otherwise.

### Feature Selection

We initially extracted over 350 million 31-mers extracted as features, and removed all those that occurred less than 10 times or more than 3,000 times. This filtering was done since these features are unlikely to be informative for classification. Next, we divided the isolates into testing and training sets and performed a chi-square test on the training set for each feature with respect to the resistance/susceptibility labels, on every antibiotic. All features were then ranked by the chi-square’s p-value. Although the chi-square test is univariate and does not recognize correlated features (subject to confounding), it is helpful to guide iterative feature selection. We further mitigated the shortcomings of the chi-square test ranking by employing incremental feature sets, shrinkage in LR, and the nonlinear RF classifier.

To operationalize the incremental feature selection, we fitted independent models with increasing numbers of features. Specifically, we considered feature sets consisting of the 2^i^ top 31-mers, based on the p-value ranking, for all integers *i* ∈ [0, 19] (i.e., from 1 to 524,288 features). In other words, we first considered a model with the top 31-mer, then considered another with the top two 31-mers, then another with the top four, eight, sixteen, thirty-two, and so forth. We continued to double the number of input features until the performance of the model did not improve over three consecutive times. This iterative approach allowed us to identify the optimal number of features maximizing the performance of the model while avoiding unnecessary complexity.

### Model Selection and Training

We trained both LR with LASSO shrinkage, and RF models for each feature subset and for each antibiotic drug, i.e., a total of 16 LR models and 16 RF models. LR and RF were chosen because they complement each other in terms of complexity, performance, and interpretability trade-offs [3]. LR is a parametric algorithm that models the probability of an outcome (i.e., being resistant to an antimicrobial) as a linear function of input features incorporated into a logistic function. In the absence of causal assumptions and/or correlation structures, the coefficients of LR are relatively simple to interpret as weights contributing to increased or decreased likelihood of the outcome. However, LR struggles with nonlinear relationships, unless they are specifically encoded as additional input variables. In contrast, RF is fully non-parametric and nonlinear. RF combines multiple decision trees, each trained on a bootstrapped subset of the data and on randomly selected feature subsets, to make class predictions through voting. Feature importance in RF can be determined in multiple ways; one is to look at the average reduction in impurity across trees. Due to its nonlinear property, RF is more flexible and capable of capturing interactions between features that the LR cannot directly handle. RF exhibits higher complexity due to multiple decision trees, which could be prone to over-fitting, but the problem is mitigated through bootstrapping and random feature selection. RF generalizes well and often delivers strong performance across tabular datasets in computational biology, even when compared to deep learning [16].

### Cross-Validation and Performance Evaluation

Cross-validation is a fundamental technique used in machine learning and statistical analysis to assess the performance of predictive models while mitigating issues related to overfitting and data variance. The core idea behind cross-validation is to divide the data into multiple subsets (or *folds*); iteratively train a model on all the folds but one, which is used as a test; and lastly, evaluate the average model performance. This process provides a more robust estimate of how well the model will generalize to new, unseen data. In this study, we used five-fold cross-validation. To assess the performance of our models and compare it to the previous models, we use F-1 score due to its robustness to data imbalance compared to other evaluation metrics. Here, a sample is considered positive for a class if it is resistant to that class.

## SUPPLEMENT

**Table 2:**
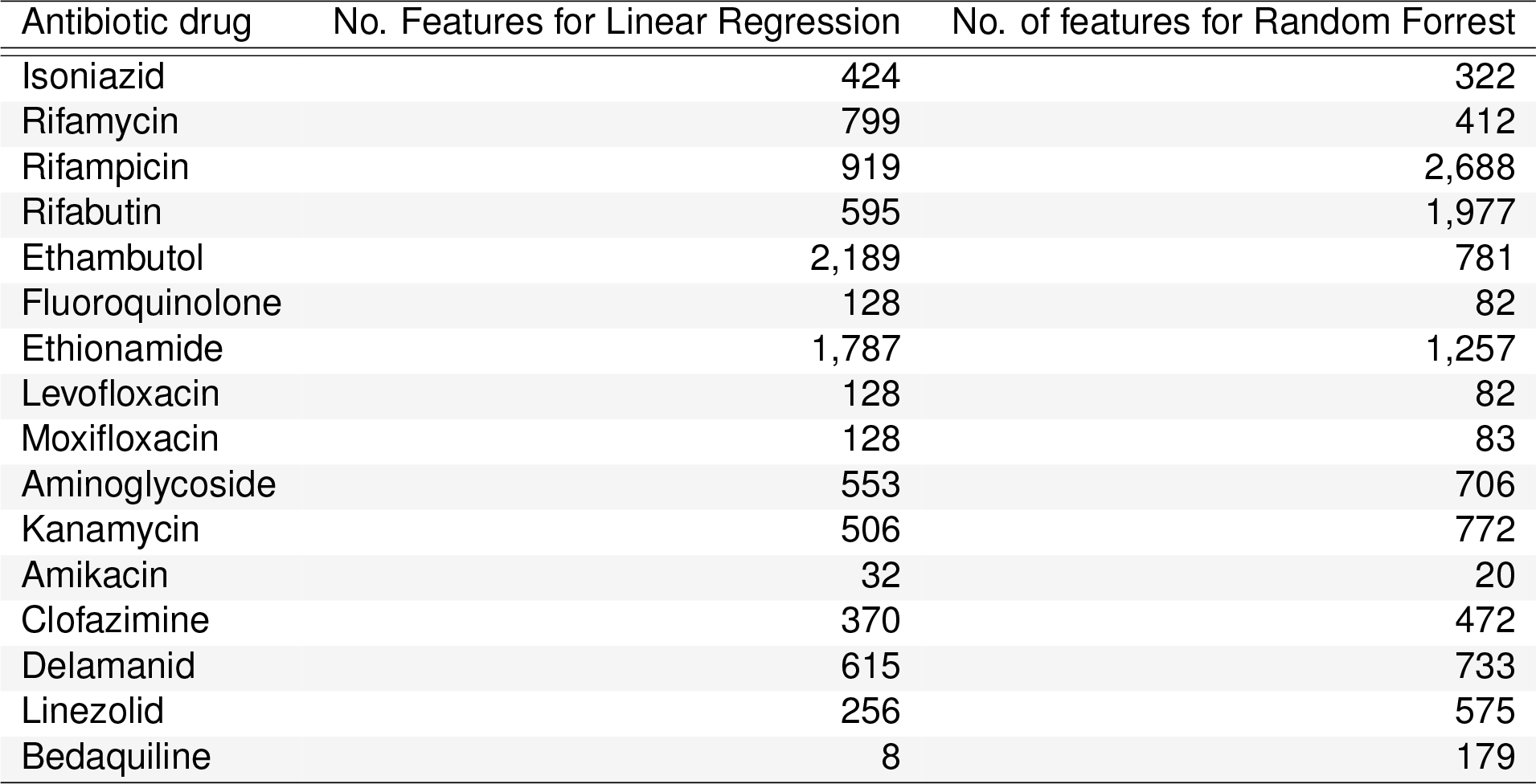
The number of features (31-mers) for each antibiotic drug is needed to optimize the F1-score of the LR and RF models.

**Table 3:** Proposed epidemiological cut-off values (ECOFF/ECVs) and suggested borderline minimum inhibitory concentrations (MICs) for three compounds [9, 33].

| Antibiotic Drug | ECOFF/ECV $\text{mg L}^{-1}$ | Borderline MIC $\text{mg L}^{-1}$ |
| --- | --- | --- |
| Isoniazid | 0.10 | 0.2, 0.4 |
| Rifampicin | 0.50 |  |
| Rifabutin | 0.12 |  |
| Ethambutol | 4.00 | 4.00 |
| Ethionamide | 4.00 | 4.00 |
| Levofloxacin | 1.00 |  |
| Moxifloxacin | 1.00 |  |
| Kanamycin | 4.00 |  |
| Amikacin | 1.00 |  |
| Clofazimine | 0.25 |  |
| Delamanid | 0.12 |  |
| Linezolid | 1.00 |  |
| Bedaquiline | 0.25 |  |

**Table 4:** Classes of resistance found in the MEGARes database using BWA for all antibiotic drugs where the balance of the data is at least 15%. The bold genes in the table are uniquely found for that specific class of resistance.

| Antibiotic drug | MEGARes Resistance Class | MEGARes Accession |
| --- | --- | --- |
| Isoniazid | MTB-specific | MEG_2710, MEG_2711, MEG_3446, MEG_8171 |
|  | Rifampin | MEG_6090, MEG_6134, MEG_7259 |
|  | Aminoglycoside | MEG_6144 |
|  | FQS | MEG_3237 |
| RIA | MTB-specific | MEG_2710, MEG_2711, MEG_3446, MEG_8171 |
|  | Rifampin | MEG_6090, MEG_6134, MEG_7259 |
|  | AMG | MEG_6144, MEG_8680 |
|  | FQS | MEG_3237 |
| Rifampicin | MTB-specific | MEG_2710, MEG_2711, MEG_3446, **MEG_8079**, MEG_8171, |
|  | Rifampin | MEG_6090, MEG_6134, MEG_7259 |
|  | AMG | MEG_3, MEG_11, MEG_6144, MEG_8680 |
|  | FQS | MEG_3237 |
| Rifabutin | MTB-specific | MEG_2710, MEG_2711, MEG_3446, MEG_8171 |
|  | Rifampin | MEG_6090, MEG_6134, MEG_7259, MEG_8675 |
|  | AMG | MEG_3, MEG_6144 |
|  | FQS | **MEG_3180**, MEG_3237 |
| Ethambutol | MTB-specific | MEG_2710, MEG_2711, **MEG_2712**, MEG_3446, MEG_8171, |
|  | Rifampin | MEG_6090, MEG_6134, MEG_7259, MEG_8675 |
|  | Aminoglycoside | MEG_11, MEG_12, MEG_6144 |
|  | FQS | MEG_3237 |
|  | Copper Resistance | **MEG_2653, MEG_2654** |
| FQS | FQS | MEG_3237 |
| Levofloxacin | FQS | MEG_3237 |
| Moxifloxacin | FQS | MEG_3237 |
| Ethionamide | MTB-specific | MEG_2710, MEG_2711, MEG_3446, **MEG_7973**, MEG_8171 |
|  | Rifampin | MEG_6090, MEG_6134, MEG_7259, MEG_8675 |
|  | FQS | MEG_3237 |
|  | AMG | MEG_3, MEG_9, MEG_11, MEG_6144, **MEG_6145**, MEG_8078, MEG_8680 |
|  | Cationic AMR peptides | MEG_1490 |

**Table 5:** Mechanisms of resistance found in the MEGARes database using BWA for all antibiotic drugs where the balance of the data is at most 11%. The bold genes in the table are uniquely found for that specific class of resistance.

| Antibiotic drug | MEGARes Resistance Class | MEGARes Accession |
| --- | --- | --- |
| AMG | Aminoglycoside | MEG_3, MEG_5, MEG_9, MEG_11, MEG_12, MEG_8078 |
|  | Rifampin | MEG_6090 , MEG_6134 |
|  | MTB-specific | MEG_3446 |
| Kanamycin | Aminoglycoside | MEG_3, MEG_5, MEG_9, MEG_11, MEG_8078 |
|  | Rifampin | MEG_6090 , MEG_6134 |
|  | MTB-specific | MEG_3446 |
|  | FQS | MEG_3237 |
| Amikacin | Aminoglycoside | MEG_5, MEG_9, MEG_11 |
| Delamanid | Drug and Biocide Resistance | **MEG_3759** |
|  | Multi-drug Resistance | **MEG_4078** |
| Linezolid | Oxazolidinone | **MEG_8670** |
|  | Cationic AMR Peptides | MEG_1490 |

## Footnotes

1 Kuang et al. [21] do not provide a trained classifier. All methods were ran with their default settings. PhyReSE is only provided as a web-interface that takes in a single genome at once and is unable to handle large batch files. GenTB failed with source code errors. The issue was posted to the GenTB GitHub: https://github.com/farhat-lab/gentb-snakemake/issues/6.

## Notes

### Competing Interest Statement

The authors have declared no competing interest.

